# The radiation of Austral teals and the evolution of flightlessness

**DOI:** 10.1101/2023.10.19.563194

**Authors:** Hanna S. Rosinger, Olga Kardialsky, Martyn Kennedy, Hamish G. Spencer, Florian M. Steiner, Birgit C. Schlick-Steiner, Nicolas J. Rawlence, Michael Knapp

## Abstract

The origin and evolution in the Southern Hemisphere of the Austral teals, consisting of the grey-teal and brown-teal species complexes, remains poorly understood due to limited molecular data. With the group containing multiple independent examples of flight loss, understanding the evolutionary history of the group is of significant interest for functional genomic studies into the evolution of flightlessness. Here we present the first whole mitogenome-based phylogeny of the Austral teals. We show that the group diverged from a common ancestor with mallards in the late Miocene and soon after radiated into the brown-teal and grey-teal lineages, as well as the widely distributed pintails and green-winged ducks. The brown-teal species complex, which includes the volant brown and Chatham Island teals as well as the flightless, sub-Antarctic Auckland and Campbell Island teals, radiated within the past 0.9 - 2.2 million years. The divergence of the extinct Chatham Island teal, and the stepping-stone colonisation of the Auckland and Campbell Islands occurred from mainland New Zealand. Morphological changes towards flightlessness are also present in the volant brown teal on mainland New Zealand, suggesting that this group was on the pathway to flightlessess, which accelerated in some insular island lineages.

## INTRODUCTION

The Austral teals (*Anas* spp.) are a group of dabbling ducks endemic to Madagascar, Australia, New Zealand, and its sub-Antarctic islands. Consisting of the Madagascar teal, the grey-teal species complex with representatives in Australia and New Zealand, and the brown-teal species complex from New Zealand and the sub-Antarctic islands (Figure 1), the origin and evolution of the Austral teals is poorly resolved. With only data from a small percentage of the mitochondrial genome available for phylogenetic analysis, this uncertainty encompasses all taxonomic levels in the group, from their relationship to other dabbling ducks such as green-winged teals (*Anas* spp.), pintails (*Anas* spp.) and mallards (*Anas platyrhynchos* Linnaeus, 1758), to the relationships within the brown-teal species complex, in particular.

**Figure One.**
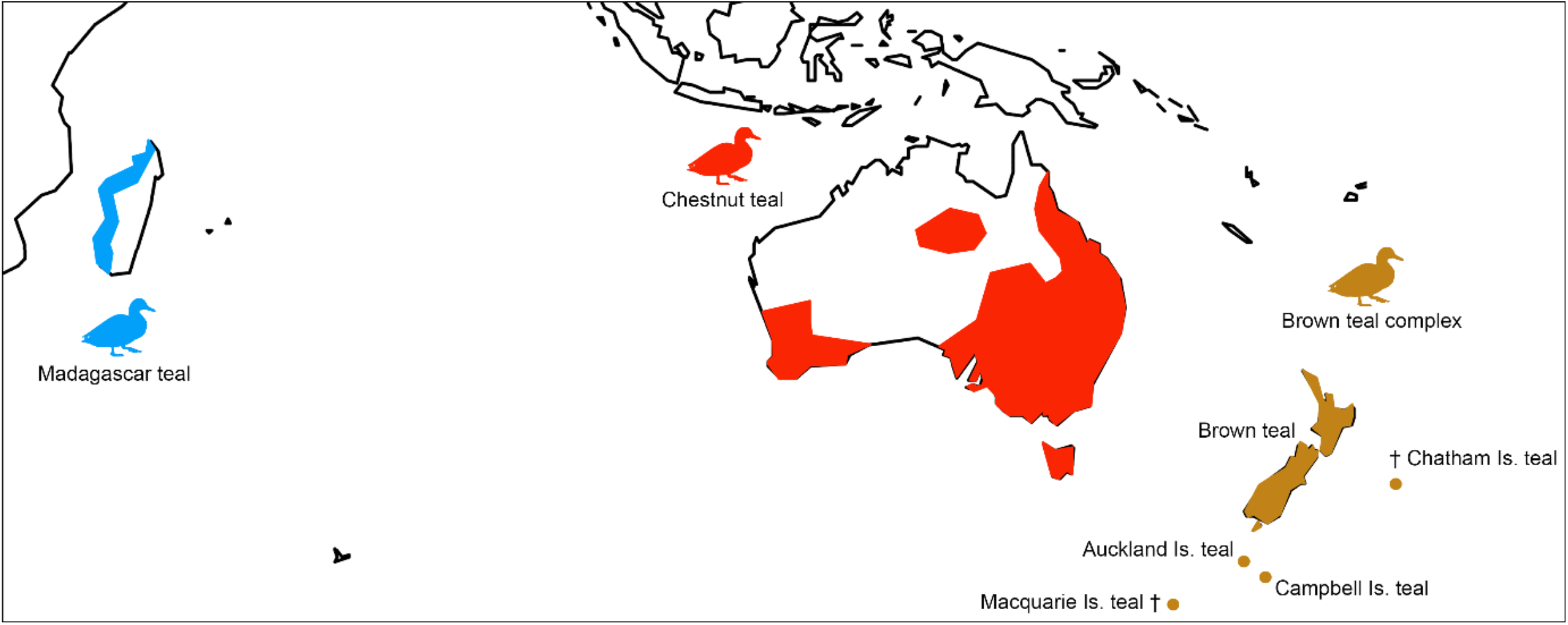
Distribution of Austral teals (*Anas* spp.) in the Southern Hemisphere including the Madagascar teal (*A. bernieri*), the chestnut teal (*A. castanea*) from Australia, and the brown-teal complex from the New Zealand region. This complex comprises the brown teal (*A. chlorotis*) from mainland New Zealand, the extinct (†) Chatham Island teal (*A. chathamica*), Auckland Island teal (*A. aucklandica*), Campbell Island teal (*A. nesiotis*), and the extinct Macquarie Island teal (*Anas* spp. indet.). The distribution of the grey teal (*A. gracilis*) from Australasia (Australia, New Guinea, New Caledonia and New Zealand) is not shown due to the overlap with chestnut and brown teal. Distributions are based off the Cornell Laboratory of Ornithology Birds of the World website.

Based on the mitochondrial cyt *b* and ND2 genes, Johnson and Sorensen (1999) identified the brown-teal species complex in the New Zealand region as an early diverging sister clade to pintails, green-winged teals, mallards and the grey-teal species complex. Gonzalez *et al*. (2009), using the same data but an alternative phylogenetic reconstruction approach, identified brown-teals and grey-teals as sister clades, which together with pintails and green-winged teals formed a sister clade to the mallard.

Within the Austral teals, the brown-teal species complex has been of special interest as it includes examples of flightlessness. The New Zealand region is home to one extinct and three living named endemic *Anas* species (Figure 1): the extinct Chatham Island teal, *A. chathamica* (Oliver, 1955); mainland brown teal, *A. chlorotis* G. R. Gray, 1845; Auckland Island teal, *A. aucklandica* (G. R. Gray, 1849); and Campbell Island teal, *A. nesiotis* (J. H. Fleming, 1935). The taxonomic status of extinct ‘brown teal’ on the Chatham Islands (nominally *A. chlorotis*) and the sub-Antarctic Macquarie Island (*Anas* sp.) is currently undetermined.

The brown teal is the only fully volant living species within the group. Just ∼2500 individuals of this species are left, with populations on Stewart Island now extinct (Williams, 2013) and on the South Island functionally extinct due to hybridization with mallards (Birdlife International, 2015). The Campbell Island teal is fully flightless (Livezey, 1990), whereas the smaller Auckland Islands teal is able to fly, but is not capable of sustained flight. It has been observed to be a weak flyer, capable only of short flights, with feet and wings splashing across the water, flapping wings to jump onto shoreline ledges and climbing stepping stones (Buller, 1905; Weller, 1975). The Chatham Island teal is the largest of the brown-teals and was formally described in a separate genus (*Pachyanas*) based on its morphological distinctiveness (Oliver, 1955). However, recent palaeogenetic research has shown it is the earliest diverging member of the brown-teal radiation (Mitchell *et al*., 2014). Further, in spite of suggestions that it was flightless (Millener, 1999), analysis of wing osteology shows that it was capable of flight (Mitchell *et al*., 2014).

There have been several attempts to resolve the phylogenetic relationships and colonization history of the brown-teal radiation using comparative morphology (Livezey, 1990; Livezey, 1991), allozymes (Daugherty *et al*., 1999), and partial mitochondrial DNA sequence data (Young *et al*., 1997; Johnson & Sorenson, 1998; Johnson & Sorenson, 1999; Kennedy & Spencer, 2000; Mitchell *et al*., 2014). The majority of studies suggest the Campbell Island teal and Auckland Island teal are sister taxa, most closely related to the brown teal (Livezey, 1991; Young *et al*., 1997; Johnson & Sorenson, 1998; Johnson & Sorenson, 1999; Mitchell *et al*., 2014). Alternatively, Kennedy and Spencer (2000) suggested that the Campbell Island teal and brown teal are sister taxa to the exclusion of the Auckland Island teal, while Daugherty *et al*. (1999) suggested the brown teal and Auckland Island teal are sister. There is also phylogenetic disagreement over the closest relatives of the brown-teal radiation – with either the grey-teal complex in Australia (Johnson & Sorenson, 1998; Johnson & Sorenson, 1999) or the Madagascar teal (Mitchell *et al*., 2014) argued to be the sister taxa of brown-teals.

The unresolved nature of the phylogenetic relationships has implications for our understanding of the evolution of the Austral teals in general, and for the colonization history of New Zealand and outlying islands, as well as the evolution of flightlessness in the brown-teal radiation. This study presents the first whole mitogenome approach to resolve the phylogeny and evolutionary history of Austral teals.

## MATERIALS AND METHODS

### Specimens

Tissue (n = 6) and feather (n = 1) samples, or genomic DNA extracts (n = 12), were sourced from the Department of Zoology (University of Otago), Auckland Museum, and the Durrell Wildlife Conservation Trust in Jersey, including: brown teal (n = 4); Auckland Island teal (n = 2); Campbell Island teal (n = 2); Madagascar teal, *Anas bernieri* (Hartlaub, 1860) (n = 4); grey teal, *Anas gracilis* Buller, 1869 (n = 2); chestnut teal, *Anas castanea* (Eyton, 1838) (n = 2); grey duck, *Anas superciliosa* Gmelin, 1789 (n = 2); and mallard (n = 1) (Table S1).

### DNA extraction

Existing DNA extracts were previously extracted by Kennedy and Spencer (2000) following a modified phenol chloroform extraction protocol by Kocher *et al*. (1989). To test for DNA degradation, fragmentation was checked on a 1 % agarose gel using GelRed (Biotium) and a 1 kbp HyperLadder (Bioline). Genomic DNA extraction from new tissue samples was conducted using the MagJet gDNA kit (Thermo Fisher Scientific) with a modified protocol based on the manufactures instructions for manual genomic DNA purification (see Supplementary Information). Successful extractions were verified by visualizing DNA using gel electrophoresis as above and quantified using a NanoDrop 2000 (Thermo Fisher Scientific).

### Whole mitogenome amplification and sequencing

Four overlapping primer pairs were designed for complete mitogenome amplification (Tables S2-S3) using published complete anatid mitogenomes: mandarin duck, *Aix galericulata* (Linnaeus, 1758); falcated duck, *Mareca falcata* (Georgi, 1775); Baikal teal, *Sibirionetta formosa* (Georgi, 1775); mallard; Indian spot-billed duck, *Anas poecilorhyncha* Forster, 1781; tufted duck, *Aythya fuligula* (Linnaeus, 1758); muscovy duck, *Cairina moschata* (Linnaeus, 1758); ruddy shelduck, *Tadorna ferruginea* (Pallas, 1764); common shelduck, *Tadorna tadorna* (Linnaeus, 1758); black swan, *Cygnus atratus* (Latham, 1790); and the lesser whistling-duck, *Dendrocygna javanica* (Horsfield, 1821). Mitogenome sequences were downloaded from Genbank and aligned in BioEdit (Version 7.2.5; Hall, 1999). The four primer pairs (F16 – R8, F3 - R10, F5043 - R3, F8 - R12734) amplified fragments with lengths ranging from 4 kb to over 5 kb (F5043 - R3: 3757 bp; F3 - R10: 4627 bp; F8 - R12734: 4854 bp; F16 – R8: 5346 bp).

Long-range PCR was conducted using a KAPA LongRange HotStart Kit (KAPA Biosystems) in 25 µl volumes with 24 µl Master-mix (ultra-pure MilliQ water, 1 x KAPA LongRange Buffer, 1.75 mM MgCl_2_, 0.3 mM dNTPs, 0.5 mM each primer, 0.025 units DNA Polymerase) and 1 µl DNA. PCR thermocycling conditions followed the KAPA LongRange HotStart DNA Polymerase kit protocol for amplification of long targets and/or low concentrations of template DNA (Table S4). A negative control was included in each PCR run to test for contamination. PCR amplification was verified by visualizing on a 1 % agarose gel with GelRed (Biotium) and 1 kb Hyperladder (Bioline). PCR products were purified using Mag-Bind RXNPurePlus beads (OMEGA Biotek) (see Supplementary Information for preparation of magnetic bead size selection buffer) following the manufactures instructions with the following modification: 1:1 volume of magnet beats, 85 % ethanol wash, 30 μl elution buffer. DNA concentrations were quantified using NanoDrop 2000 (Thermo Fisher Scientific).

PCR products for each sample were pooled at equimolar ratios and sonicated using a Picorupter (Diagenode) to shear the DNA into small fragments (seven cycles of 15 sec on and 45 sec off). Successful fragmentation was checked on a 2 % agarose gel. After sonication and in between each of the steps in double-stranded DNA library construction, DNA purification was conducted as above using Mag-Bind RXN Pure Plus beads.

Indexed double-stranded barcoded libraries for high-throughput sequencing were prepared from the purified sheared long-range PCR products following a modified Illumina TruSeq protocol (see Supplementary Information). Quantification of libraries was conducted with a Qubit 2.0 fluorometer (Invitrogen) following the manual assay preparation instructions. Libraries were pooled at equimolar concentrations and a final concentration was determined using the Qubit 2.0 (Invitrogen). High-throughput sequencing was conducted at the Otago Genomic & Bioinformatics Facility (University of Otago) on the Illumina MiSeq platform.

### Bioinformatic and phylogenetic analysis

Mitogenomes were assembled in Geneious (v.10.1.3, Kearse *et al*., 2012) using the mallard as a reference genome (EU009397.1). The resulting mitogenomes were aligned, annotated and manually edited to exclude evident errors in BioEdit (v.7.2.5; Hall, 1999). Our phylogenetic analysis utilized complete mitogenomes supplemented with published mitogenome sequences of *Anas* spp. including two each of the Indian spot-billed duck, mallard, falcated duck, northern shoveler *Spatula clypeata* (Linnaeus, 1758), northern pintail *Anas acuta* Linnaeus, 1758, Baikal teal, Eurasian teal *Anas crecca* Linnaeus, 1758, one Chatham Island teal and outgroup mitogenomes including two each of the redhead *Aythya americana* (Eyton, 1838) and the common pochard *Aythya ferina* (Linnaeus, 1758) (see Table S2). We used the same fossil calibration points as Mitchell et al. (2014). Sequences were aligned in BioEdit using the default settings (v7.2.5; Hall, 1999) and checked by eye.

Phylogenetic analyses were conducted with an alignment that was partitioned into individual genes, tRNAs and rRNAs. The control region/D-loop was excluded from the analyses due to ambiguous alignment. Genes were further partitioned into 1st, 2nd, and 3rd codon positions. The best fitting models of nucleotide substitution were identified using the Akaike Information Criterion as implemented in jModeltest2 (Darriba *et al*., 2012). Downstream phylogenetic and molecular clock analyses were run with the following substitution models: rRNA, TPM1uf +I; tRNA, GTR+I; genes (all codon positions), TIM2+I+G.

We used BEAST (v2.7.4; Bouckaert *et al*., 2014) for phylogenetic reconstructions and molecular clock analyses. We followed Mitchell *et al*. (2014) to fossil calibrate our dataset under a Bayesian framework. We modelled the stem age of *Anas* using a lognormal prior distribution (M=1, S=1, offset 11.2) so that 95 % of the prior probability fell between 11.6 and 30.5 million years ago (Mya). All other group priors had no prior settings. We applied a strict molecular clock and an optimised relaxed molecular clock using the Yule prior and birth-death priors, respectively, with a 10,000,000 generation chain. Results were visualized in Tracer (v.1.5.0) to evaluate whether they adhered to a strict or a relaxed molecular clock model and which tree prior best described the data. The different settings of (1) strict and relaxed molecular clock with the Yule prior and (2) strict and relaxed molecular clock with the birth-death prior were then compared using Bayes factors and their suitability was interpreted following Kass and Raftery (1995). After Bayes factor analysis, a relaxed clock (three-independent 20,000,000 generation chains) with a Yule prior was applied to the dataset to estimate the substitution rate and divergence times. After checking for Markov Chain Monte Carlo (MCMC) convergence for each run, all three runs were combined using LogCombiner (v.2.7.4) to ensure that all parameters had reached effective sample sizes (ESS) of 200 or more. TreeAnnotator (v.2.7.4) was used to combine the independent trees with a 25% burn-in and FigTree (v1.4.4) was used to construct the chronogram.

## RESULTS

### Mitogenome sequencing

Long-range PCR amplification of the mitogenome was successful for all 19 samples. High throughput sequencing resulted in a mean depth of coverage for the mitogenome of 71x to 4225x, with the number of mapped reads ranging from 5,524 to 22,194, producing mitogenomes of ∼16,850 bp with 99.8% to 100% coverage of the mallard reference genome.

### Phylogenetic analysis

The time-calibrated Bayesian phylogenetic analysis (Figure 2) shows that the Austral teals, as well as the widely distributed pintails and green-winged ducks, form a clade that diverged from the mallards in the late Miocene, approximately 8 Mya. Approximately one million years later, the Austral teals diverged from the pintails and green-winged ducks, although this divergence is poorly resolved and is – based on our data – best considered a polytomy that consists of four major clades: 1) pintails, 2) green-winged ducks, 3) brown-teal complex, 4) grey-teal complex and Madagascar teal.

**Figure Two.**
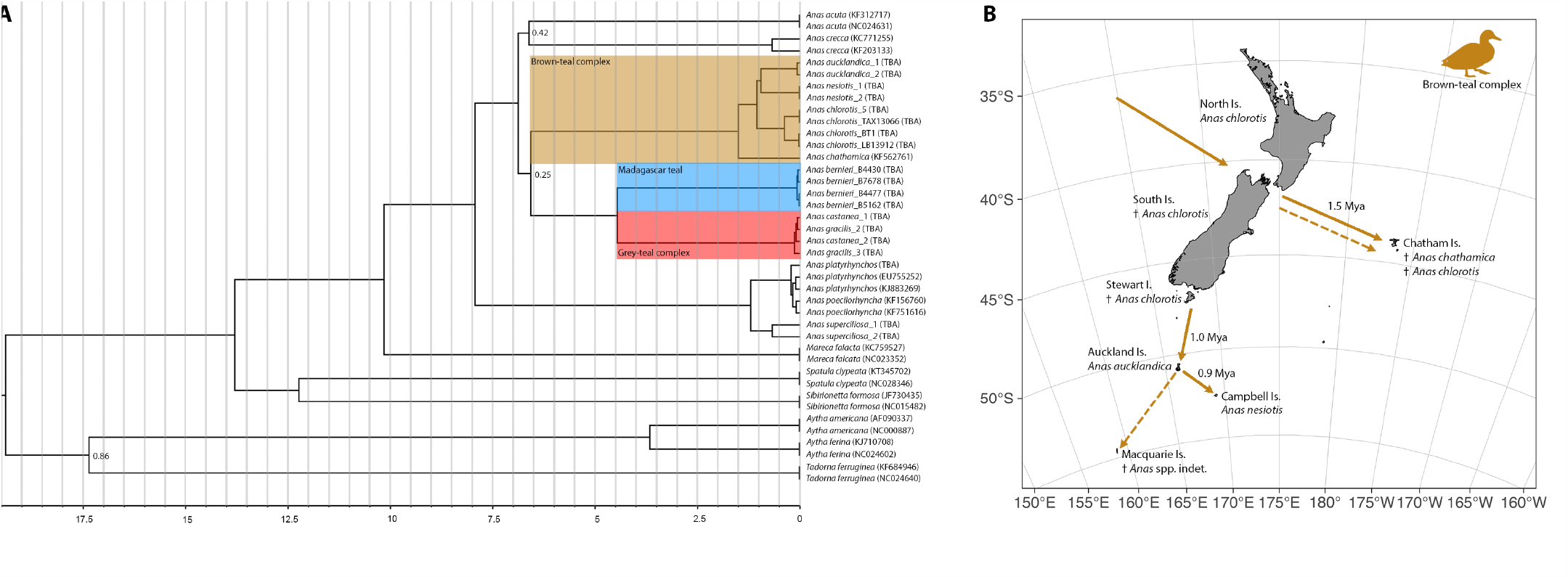
(**A**) Time-calibrated Bayesian phylogeny of Austral teal mtDNA created using BEAST. Branch lengths are proportional to time in millions of years before present. Posterior probability support values are shown by the nodes subtending the branches they belong to if less than 1.0. Divergence times and 95% highest posterior densities of node estimates are shown in Table 1. Tips are labelled with species names and GenBank accession numbers. (**B**) Schematic of colonisation routes of the New Zealand by the brown-teal complex. Divergence times are mean estimates based on our phylogenetic analysis in (A) and Table 1. Solid arrows represent known dispersal events, while dashed arrows represent inferred dispersal events.

Within the brown-teal complex, the ancestors of the Chatham Island teal diverged first (1.5 Mya, 95% HPD 0.9 – 2.2 Mya), followed by the brown teal (1.1 Mya, 95% HPD 0.7 – 1.5 Mya). Auckland Island teal and Campbell Island teal diverged from each other soon after (1 Mya, 95% HPD 0.6 – 1.4 Mya) (Table 1). The sister relationship between the Auckland Island and Campbell Island teal is highly supported (1.0 posterior probability), despite the short internodal distances between branches in the brown-teal clade.

**Table 1.**
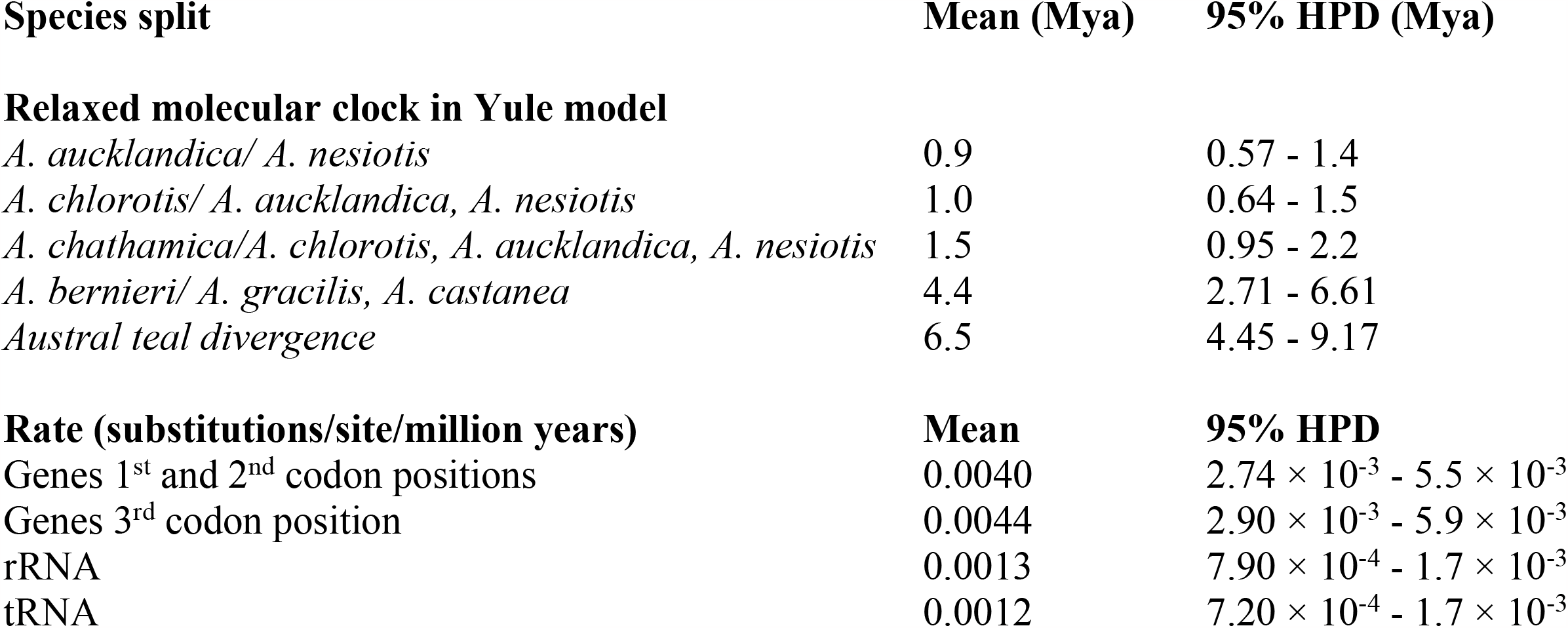
Estimated divergence times and substitution rates of Austral teals calculated in BEAST.

The Madagascar teal clade diverged from the Australian and New Zealand grey-teal complex (which occurs in Australia, New Guinea, New Caledonia and New Zealand) 4.5 Mya (95% HPD 2.7 – 6.5 Mya) (Table 1). Within the recent crown group grey-teal complex, chestnut teals and grey teals are not reciprocally monophyletic.

The estimated substitution rates (in substitutions/site/million years) were: genes 1^st^ and 2^nd^ codon positions: 0.004 (95% HPD 2.74 × 10^-3^ to 5.5 × 10^-3^); genes 3^rd^ codon position: 0.0044 (95% HPD 2.9 × 10^-3^ to 5.9 × 10^-3^); rRNA: 0.0013 (95% HPD 7.9 × 10^-4^ to 1.7 × 10^-3^); tRNA: 0.0012 (95% HPD 7.2 × 10^-4^ to 1.7 × 10^-3^).

## DISCUSSION

### Evolution of the Austral teals

The Austral teals are a group of dabbling ducks endemic to the Southern Hemisphere, which includes the grey-teal and brown-teal species complexes, as well as the Madagascan teal (Figure 1). Their origin and evolution, however, has remained poorly resolved. Our phylogenetic analysis of whole mitogenomes (Figure 2, Table 1) shows Austral teals diverged during the late Miocene from the common ancestor of the widespread Northern Hemisphere mallard. However, we were unable to determine whether there was (1) a single pulse of dispersal into the Southern Hemisphere and then subsequent divergence into the Madagascan teal, grey-teal and brown-teal complexes, or (2) two independent dispersal events, one leading to the brown-teals and the other to the Madagascan and grey teals. The rapid radiation between the pintail, Eurasian teal, and brown-teal species complexes, and a strongly supported clade containing the Madagascan teal and grey-teal species complex, was too fast for the evolution of informative single nucleotide polymorphisms that would have resolved the branching patterns and phylogenetic relationships.

Our results are in contrast to Mitchell *et al*. (2014), for example, who suggested the brown-teal radiation was closest to the Madagascan teal. Our findings strengthen the hypothesis that the Madagascan, grey and chestnut teals are closely related (see Kennedy & Spencer, 2000) (Figure 2), and that the paraphyletic nature of grey and chestnut teals in Australia (forming a single intermixed clade rather than two reciprocally monophyletic clades; Figure 2) indicates high levels of hybridization (Donne-Gousse *et al*., 2002). Future genomic research, involving nuclear data, and additional samples of other dabbling ducks like green-winged teals (e.g., Spaulding *et al*., 2023) and pintails, is therefore needed to help resolve deeper nodes in our phylogeny and the evolution of Austral teals.

### Phylogeography of brown teals

Within Austral teals, the phylogeography and evolution of flightlessness in the brown-teal complex also remains unresolved. Our analyses support previous assertions that the Auckland and Campbell Island teal are sister taxa (Young *et al*., 1997; Johnson & Sorenson, 1999; Mitchell *et al*., 2014), contrary to other studies (e.g. Daugherty *et al*., 1999; Kennedy & Spencer, 2000).

Our phylogenetic analyses imply that over a short period of time brown-teal ancestors dispersed from mainland New Zealand to the Chatham Islands, and a subsequent independent stepping stone dispersal to the Auckland, and then, Campbell Islands (Figure 2). However, we cannot fully exclude the parallel colonization scenario, with the Auckland and Campbell Islands’ independently populated from mainland New Zealand. For parallel colonization, two scenarios are possible: (1) the Auckland and Campbell Islands’ founding populations originated from the same ancestral population on mainland New Zealand, which later became extinct (Mitchell *et al*., 2014); and (2) the divergence occurred on mainland New Zealand in combination with subsequent dispersal to each of the sub-Antarctic islands and their dying out on the mainland. Scenario two seems less parsimonious, however. Independent colonisation events of the Chatham Islands, mainland New Zealand, and the Auckland and Campbell Islands (as suggested by Williams *et al*., 1991) from Australia can be ruled out given the monophyly of the brown-teal radiation.

### Age and evolution of flightlessness in brown teals

An upper limit for the age of a (flightless) island species is the age of island formation, although the actual age can be much smaller (Slikas *et al*., 2002). The Auckland and Campbell Islands are all of late Miocene/early Pliocene volcanic origin (Adams *et al*., 2008; Scott & Turnbull, 2019), and therefore considerably older than the estimated divergence times in the brown-teal species complex. Flightlessness in this complex likely evolved within the past 1.4 million years during the Pleistocene. While this estimate provides a maximum timeframe for the loss of flight, it does not allow for any conclusions about the actual time it took for the sub-Antarctic teals to lose their ability to fly. Loss of flight can be rapid and has frequently occurred in other bird lineages. The extinct New Zealand Finsch’s duck *Chenonetta finschi* (Van Beneden, 1875), is thought to have become flightless in just 9000 years (Worthy, 1998). In the genus *Tachyeres*, three species became flightless within the past 1.4 – 0.015 million years (Fulton *et al*., 2012): the Fuegian steamer duck *T. pteneres* (Forster, 1844), the Chubut steamer duck *T. leucocephalus* Humphrey & Thompson, 1981, and the Falkland steamer duck *T. brachypterus* (Latham, 1790). Flightlessness evolved independently in a number of *Porzana* rails within the last 0.125 – 0.5 million years (Slikas *et al*., 2002). On the sub-Antarctic islands, numerous species became (near) flightless including the Auckland Island rail *Lewinia muelleri* (Rothschild, 1893), Chatham Island coot *Fulica chathamensis* Forbes, 1892, Dieffenbach’s rail *Gallirallus dieffenbachia* (G. R. Gray, 1843), and Chatham Island rail *Cabalus modestus* (Hutton, 1872).

Flightlessness in the brown-teal species complex likely evolved as an adaptation to new environmental conditions (e.g., isolated islands, extreme prevailing westerly winds) and a lack of predators (James & Olson, 1983; Livezey, 1990; Williams, 1995; Trewick, 1997; Worthy, 1988; Kennedy & Spencer, 2000) but probably not for energy conservation given flight is energetically expensive (McNabb, 1994; McNabb, 2003).

It is possible that the insular island lineages within the brown-teal species complex were pre-adapted towards flightlessness (i.e., a shift towards flightlessness was already occurring), given morphological characteristics in the brown teal compared to the grey teal such as shortening of the wing bones and coracoid, relative wing length, wing loading, reduced strength in pectoral girdle elements and reduced depth of sternal carina (Livezey, 1990; Worthy, 1988). However, these changes could also result from adaptive responses to selective pressures such as a terrestrial lifestyle through arrested development in adults (Lack, 1970; Weller, 1975; Weller, 1980; Livezey, 1990; Trewick, 1997).

## CONCLUSION

The origin and evolution of the Austral teals has long interested scientists, especially given the incidence of flightlessness within this group. Our data suggests a radiation the Austral teals in the late Miocene. Within the brown-teal species complex, we were able to infer a basal divergence of the extinct Chatham Island teal, and the likely stepping-stone colonization of the sub-Antarctic islands from mainland New Zealand within the past 1.4 million years. Our data therefore provide a phylogenetic and temporal framework for the evolution of the group. These results can serve as a foundation for more in-depth, functional genomic studies focussing on the molecular basis of flightlessness in insular island lineages.

## ACKNOWLEDGEMENTS

Thank you to Glyn Young (Durrell Wildlife Conservation Trust), Andrew Excley, Murray Williams, Brent Evans and Charles Daugherty for providing specimens for genetic analysis. Māori are kaitiaki (guardians) of the fauna and flora of Aotearoa New Zealand, with which they are interconnected through shared whakapapa (genealogy). We would like to thank the Ngāi Tahu Murihiku Kaitiaki Rōpū Committee for their kind support of this study.

## DATA AVAILABILITY

The data that support the findings of this study are available from … [DOIs and/or accession numbers TBA].

## SUPPLEMENTARY INFORMATION

### Genomic DNA extraction

1. Tissue preparation
  a. Feathers: The tip of each feather, which was covered with blood and skin cells, was cut off and homogenized with a sterile razor blade. The tissue was placed into a 2.0 ml micro centrifuge tube, which was prefilled with 250 μl digestion buffer and 20 μl proteinase K. To obtain a uniform suspension the tube was mixed by pipetting. Samples were placed into a rotating incubating oven at 56°C for 24 hours. After incubation, an additional 50 μl digestion buffer was added. Tubes were spun down for 20 sec at 10,000 rpm. 300 μl of the resulting lysate and 300 μl lysis buffer was added to a new tube and vortexed for 5-10 sec.
  b. Tissues: A small piece of the muscle was removed with a sterile razor blade and placed in a 2.0 ml micro centrifuge tube which was prefilled with 200 μl digestion buffer and 20 μl proteinase K. Samples were placed into a rotating incubating oven at 56°C for 24 hours. Tubes were spun down for 20 sec at 10,000 rpm. 300 μl of lysis buffer was added to each tube and vortexed for 5-10 sec.
2. For tissue and feather samples, the lysate was transferred to new tubes prefilled with 400 μl isopropanol and 25 μl magnetic beads, and resuspended well by vortexing for 5-10 sec.
3. Tubes were placed on a magnetic rack for 3 min to let the magnetic beads collect at the magnet. The supernatant was removed by pipette and discarded.
4. Tubes were removed from the magnetic rack and 800 μl wash buffer 1 was added to each sample. The magnetic beads were resuspended by vortexing 5-10 sec. Tubes were placed on the magnetic rack to let the magnetic beads collect at the magnet for 2-3 min. The supernatant was removed by pipette and discarded.
5. Tubes were removed from the magnetic rack and 800 μl wash buffer 2 was added to each sample. The magnetic beads were resuspended by vortexing 5-10 sec. Tubes were placed on the magnetic rack to let the magnetic beads collect at the magnet for 2-3 min. The supernatant was removed by pipette and discarded.
6. Step (5) was repeated.
7. To make sure that any remaining supernatant was completely removed, tubes were spun down for 20 sec at 10,000 rpm, placed on a magnetic rack for 3 min, and the supernatant was removed by pipette and discarded. Tubes were left open to air dry at room temperature for 10 min.
8. Tubes were removed from the magnetic rack and 100-150 μl elution buffer was added to each sample. The magnetic beads were resuspended by vortexing 5-10 sec. The tubes were incubated in a thermomixer at 72°C for 5 min with 2-3 times vortexing in between. The tubes were span down for 20 sec at 10,000 rpm and then placed on the magnetic rack to let the magnetic beads collect at the magnet for 2-3 min.
9. While on the magnetic rack, the eluate containing purified DNA was transferred by pipette to a new clean tube.
10. If needed, step (8) was repeated to obtain more DNA.

### Preparation of Mag-Bind RXNPurePlus magnetic bead size selection buffer (OMEGA) for DNA purification

1. 50 mL of Tris EDTA (TE) buffer (10 mM Tris-HCl, 1mM EDTA) was prepared.
2. 1 ml of vortexed Sera-Mag Speed Beads solution was added to a 1.5 ml microtube.
3. The Speed Bead solution was placed on a magnetic stand until the beads were drawn to the magnet. The supernatant was removed and discarded.
4. 1 ml TE buffer was added to the Speed Beads, which were resuspended and placed on a magnetic stand until the beads were drawn to the magnet. The supernatant was removed and discarded.
5. 1 ml TE buffer was added to the Speed Beads and resuspended.
6. 9 g PEG-8000, 10 ml 5 M NaCl, 500μl 1M Tris-HCl, and 100μl 0.5 M EDTA was added to a new 50 ml falcon tube. The tube was filled to ∼49 ml using sterile water. The tube was mixed for 3-5 minutes until the PEG-8000 was in solution.
7. 27.5 μl Tween-20 was added to the PEG-8000 solution from step (6) and mixed gently.
8. 1 ml Speed Beads from step (5) was added to the PEG-8000/Tween-20 solution.
9. The tubes were wrapped in tinfoil and stored at 4°C.

### Modified Illumina TruSeq double stranded library preparation protocol

#### Blunt end repair

Blunt end repair was conducted using 1 x Buffer Tango, 4 mM dNTPs, 1 mM ATP, 0.5 U/μl T4 PNK, 0.1 U/μl T4 Polymerase, 30 μl of sonicated fragmented long-range PCR product, and UltraPure distilled water up to 40 μl. The tubes were incubated on a thermal cycler at 12°C for 55 min followed by 25°C for 15 min. Blunt end repaired products were purified using Mag-Bind RXNPurePlus beads (OMEGA Biotek) following the manufactures instructions with the following modification: 1:1 volume of magnet beats, 85 % ethanol wash, 30 μl elution buffer.

#### A-tailing

A-tailing was conducted using NEBuffer 2 (New England Biolabs), 5.7 mM Klenow fragment (3’>5’ exo), 0.1 mM dATP, and 25 μl of purified blunt end repaired DNA. Tubes were incubated at 37°C for 30 min. A-tailed products were purified using Mag-Bind RXNPurePlus beads (OMEGA Biotek) following the manufactures instructions with the following modification: 1:1 volume of magnet beats, 85 % ethanol wash, 30 μl elution buffer.

#### Ligation of truncated P5 and P7 adapters

30 μl of eluate from A tailing was combined with 1.0 μl each of double-stranded indexing adapters P5 and P7. An adapter ligation mix was created with 1 x T4 ligase buffer, 5 % PEG-4000, 1 mM T4 ligase, and UltraPure distilled water up to 16 μl. The A tailing eluate/adapter mix was added to 10 μl ligation mix. Tubes were incubated at 22°C for 1 hour in a thermocycler. Adapter ligated products were purified using Mag-Bind RXNPurePlus beads (OMEGA Biotek) following the manufactures instructions with the following modification: 1:1 volume of magnet beats, 85 % ethanol wash, 30 μl elution buffer.

#### Adapter extension and barcoding PCR

We follow the manufactures instructions for the KAPA HiFi HotStart PCR kit: briefly, the master mix per sample included 1x KAPA HiFi buffer, 0.3 mM dNTPs, 0.3 μM of each indexing primer, 0.1 U KAPA HiFi HotStart DNA polymerase, and UltraPure distilled water up to 50 μL. Reactions were incubated under the following thermocycling conditions: 94°C 3 min, 15 x [94°C 25 sec, 60°C 15 sec, 72°C 15 sec], 72°C 15 min. Barcoded products were purified using Mag-Bind RXNPurePlus beads (OMEGA Biotek) following the manufactures instructions with the following modification: 1:1 volume of magnet beats, 85 % ethanol wash, 30 μl elution buffer.

**Table S1.** Austral teal specimens used in this study to generate mitogenomes.

| **Taxonomic name** | **Common name** | **Sample ID** | **Sample location** | **Sample date** | **Collector** | **Sample type** |
| --- | --- | --- | --- | --- | --- | --- |
| *Anas aucklandica* | Auckland Island teal | AIT1 | Auckland Island | 1999 | Daugherty *et al.*, (1999)* | DNA |
| *Anas aucklandica* | Auckland Island teal | AIT2 | Auckland Island | 1999 | Daugherty *et al.*, (1999)* | DNA |
| *Anas bernieri* | Madagascar teal | B4477 | Durrell Wildlife Trust Jersey, UK | 2017 | Glyn Young | Tissue |
| *Anas bernieri* | Madagascar teal | B7678 | Durrell Wildlife Trust Jersey, UK | 2017 | Glyn Young | Tissue |
| *Anas bernieri* | Madagascar teal | B4430 | Durrell Wildlife Trust Jersey, UK | 2017 | Glyn Young | Tissue |
| *Anas bernieri* | Madagascar teal | B5162 | Durrell Wildlife Trust Jersey, UK | 2017 | Glyn Young | Tissue |
| *Anas castanea* | Chestnut teal | CT1 | Australia | 1999 | Daugherty *et al.*, (1999)* | DNA |
| *Anas castanea* | Chestnut teal | CT2 | Australia | 1999 | Daugherty *et al.*, (1999)* | DNA |
| *Anas chlorotis* | Brown teal | TAX13066 | Okiwi, Great Barrier Island | - | - | Tissue |
| *Anas chlorotis* | Brown teal | LB13912 | Mimiwhangata Bay | 2003 | Andrew Excley | Tissue |
| *Anas chlorotis* | Brown teal | BT1 | Northland | 1996 | Murray Williams | DNA |
| *Anas chlorotis* | Brown teal | BT5 | Great Barrier Island | 1999 | Daugherty *et al.*, (1999)* | Tissue |
| *Anas gracilis* | Grey teal | GT2 | Manawatu | 1990 | - | DNA |
| *Anas gracilis* | Grey teal | GT3 | Manawatu | 1990 | - | DNA |
| *Anas nesiotis* | Campbell Island teal | CIT1 | Campbell Island | 1999 | Daugherty *et al.*, (1999)* | DNA |
| *Anas nesiotis* | Campbell Island teal | CIT2 | Campbell Island | 1999 | Daugherty *et al.*, (1999)* | DNA |
| *Anas platyrhynchos* | Mallard | Mallard | Otago | 1996 | Brent Evans | DNA |
| *Anas superciliosa* | Grey duck | Grey D1 | North Island | - | Murray Williams | DNA |
| *Anas superciliosa* | Grey duck | Grey D2 | North Island | - | Murray Williams | DNA |

\* Daugherty CH, Williams M, Hay JM. 1999. Genetic differentiation, taxonomy and conservation of Australasian teals *Anas* spp. *Bird Conservation International* 9: 29-42.

**Table S2.** Published mitogenomes used for long range PCR primer design and phylogenetic analysis.

| **Taxonomic name** | **Genbank number** |
| --- | --- |
| **Long-range PCR primer design** |  |
| *Anas platyrhynchos* | EU009397.1 |
| *Anas poecilorhyncha* | NC_022418.1 |
| *Mareca falcata* | NC_023352.1 |
| *Sibirionetta formosa* | JF730435.1 |
| *Aix galericulata* | KF437906.1 |
| *Aythya fuligula* | NC_024595.1 |
| *Cairina moschata* | NC_010965.1 |
| *Cygnus atratus* | NC_012843.1 |
| *Dendrocygna javanica* | FJ379296.1 |
| *Tadorna ferruginea* | NC_024640.1 |
| *Tadorna tadorna* | NC_024750.1 |
| **Phylogenetic analysis** |  |
| *Anas acuta* | NC_024631.1 |
| *Anas acuta* | KF312717.1 |
| *Anas chathamica* | KF562761.1 |
| *Spatula clypeata* | NC_028346.1 |
| *Spatula clypeata* | KT345702.1 |
| *Anas crecca* | KF203133.1 |
| *Anas crecca* | KC771255.1 |
| *Mareca falcata* | NC_023352.1 |
| *Mareca falcata* | KC759527.1 |
| *Sibirionetta formosa* | NC_015482.1 |
| *Sibirionetta formosa* | JF730435.1 |
| *Anas platyrhynchos* | EU755252.1 |
| *Anas platyrhynchos* | KJ883269.1 |
| *Anas poecilorhyncha* | KF751616.1 |
| *Anas poecilorhyncha* | KF156760.1 |
| *Aythya americana* | AF090337.1 |
| *Aythya americana* | NC_000877.1 |
| *Aythya ferina* | NC_024602.1 |
| *Aythya ferina* | KJ710708.1 |
| *Tadorna ferruginea* | NC_024640.1 |
| *Tadorna ferruginea* | KF684946.1 |

**Table S3.**
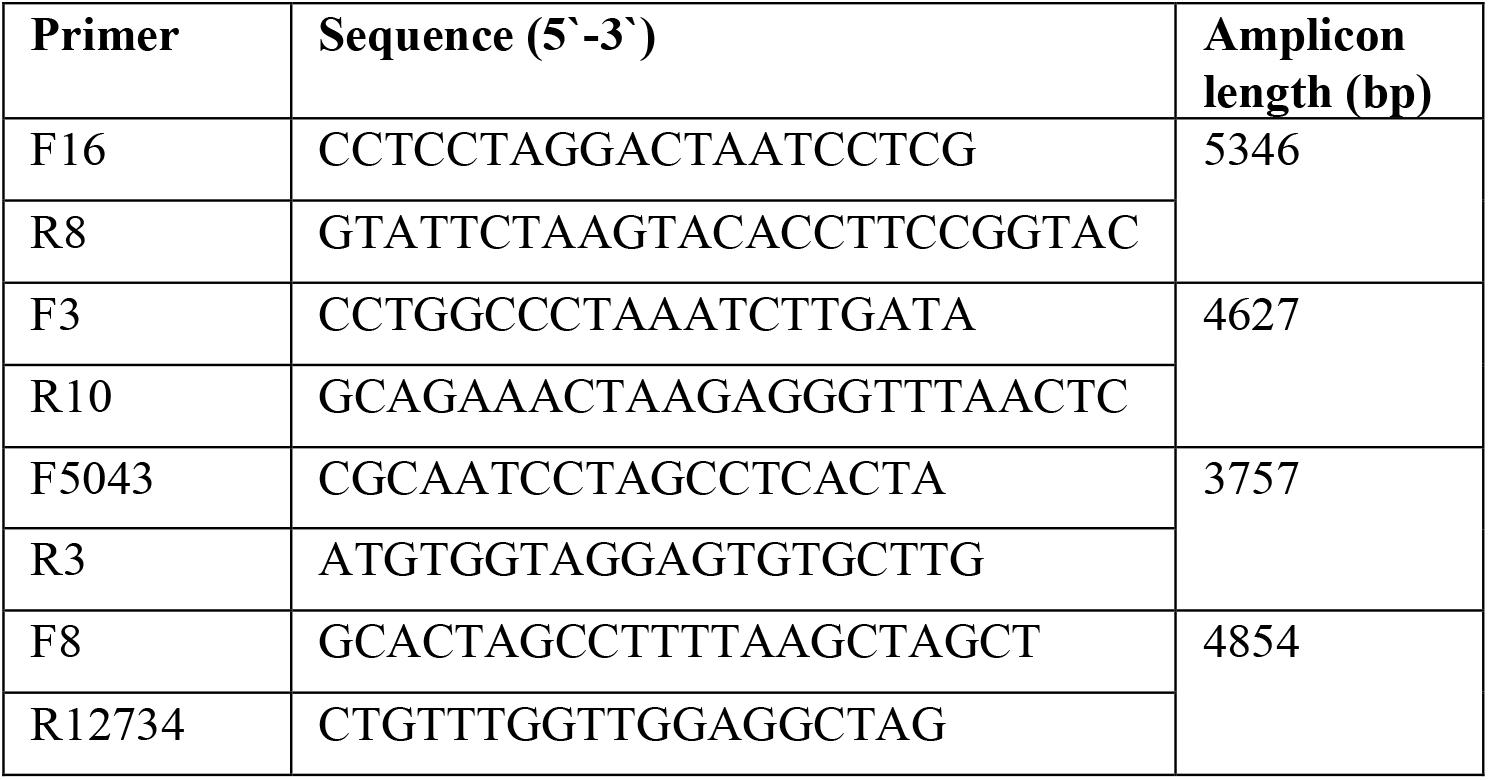
Long range PCR primers used in this study to amplify mitogenomes.

**Table S4.**
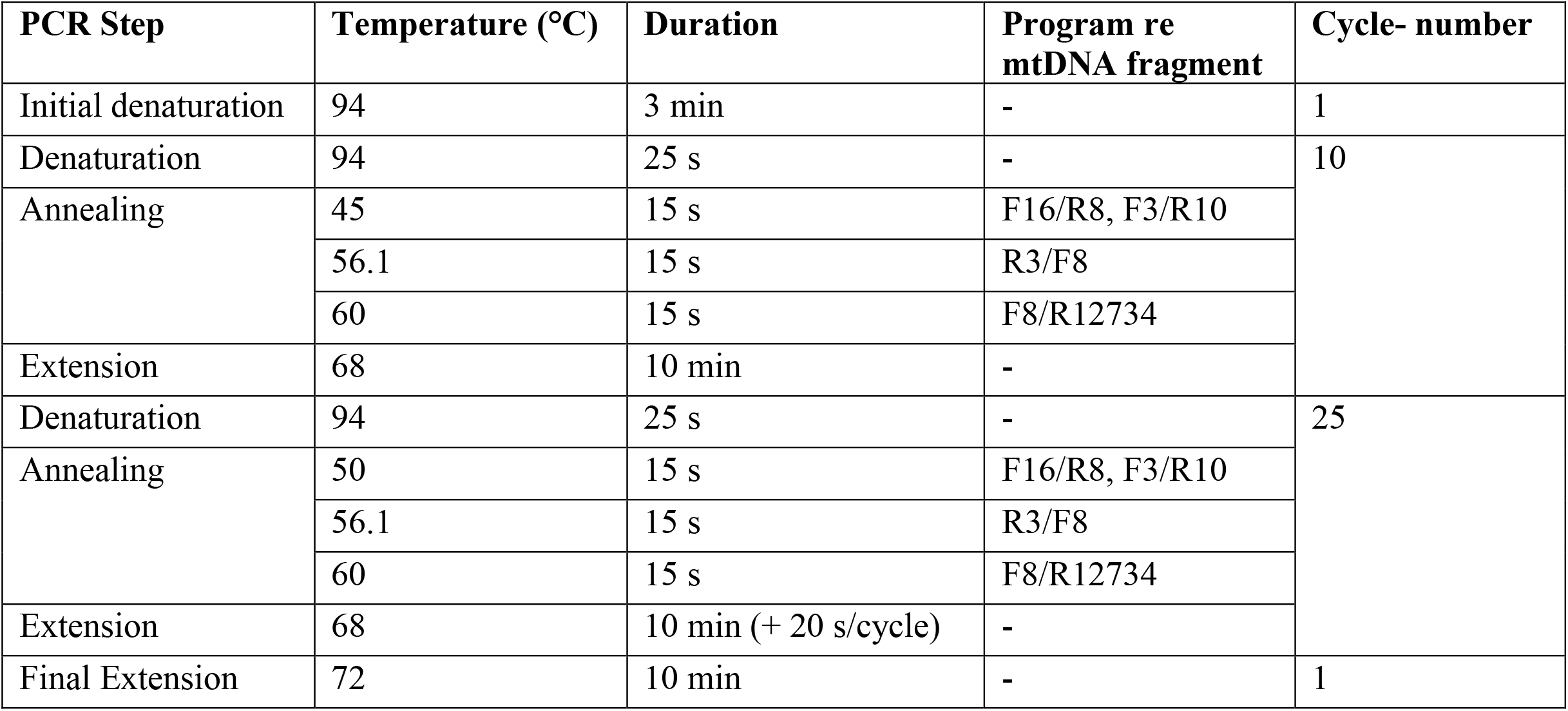
Long range PCR thermocycling conditions.

